# Delayed indel formation after Cas9 cleavage promotes mosaicism and aneuploidy in human embryos

**DOI:** 10.1101/2025.05.07.652614

**Authors:** Alex Robles, Julie Sung, Stepan Jerabek, Jishnu Talukdar, Diego Marin, Jia Xu, Nathan Treff, Dieter Egli

## Abstract

Cas9 provides a powerful tool to interrogate DNA repair and to introduce targeted genetic modifications. However, a major challenge of Cas9-based editing in human embryos is the occurrence of chromosomal abnormalities caused by Cas9 cleavage. Furthermore, mosaicism - different genetic outcomes in different cells, prevent accurate genotyping using a single embryo biopsy. Through timed analysis of editing outcomes during the first cell cycle and timed inhibition of Cas9 using AcrIIA4, we show that most edits occur at least 12 hours post Cas9 injection and therefore after the first S-phase. This timing limits the ability to achieve uniform editing across cells. We found that segmental chromosomal abnormalities and the consequential loss of heterozygosity are common at Cas9 cleavage sites throughout the genome, including at MYBPC3 and at CCR5 loci, for which this has not previously been reported. Surprisingly, inhibiting Cas9 activity 8-12 hours before mitosis does not eliminate chromosomal aneuploidies. This suggests that double-strand break (DSB) repair in human embryos is exceedingly slow, with breaks remaining unrepaired for many hours. Thus, the timing of DSB induction and repair relative to the first S-phase and the first mitosis is intrinsically limiting to preventing mosaicism and maintaining genome stability in embryonic gene editing.

## INTRODUCTION

CRISPR (clustered regularly interspaced short palindromic repeats)–Cas9 is a revolutionary gene-editing technology that uses a bacterial-derived DNA endonuclease to induce double-strand breaks (DSB) at precise locations in the genomes of model organisms and cultured human cells (1). The ability to introduce precise genetic modifications makes it a promising tool for altering genotypes, including correction of disease-causing mutations. Prior studies have attempted to use CRISPR-Cas9 to target several diseases in somatic cells, such as sickle cell disease, cystic fibrosis, vision impairment, severe combined immunodeficiency (SCID), and others. Moreover, the application of CRISPR-Cas9 in human embryos may prevent disease-causing mutations from propagating in the human germline (2).

However, the editing of the genome in the human embryo faces formidable biological obstacles, in addition to ethical and regulatory questions, with the latter having been discussed elsewhere (3). Genetic mosaicism-disparate genetic outcomes in different cells of the same embryo- is incompatible with accurate genotyping. Furthermore, the introduction of a DNA break can result in the loss of chromosome segments and whole chromosomes (4, 5). Additionally, off-target editing can occur in highly similar sequences at other locations in the genome, resulting in repaired inDels or chromosomal aneuploidy in those chromosomes. Interestingly, off-target editing appears to be temporally delayed relative to on-target editing (4, 6), increasing the risk of mosaicism and inaccurate genotyping due to divergence between trophectoderm biopsy and inner cell mass.

The timing of CRISPR-Cas9 activity plays a significant role in the uniformity of gene editing. When applied at the time of fertilization with sperm, Cas9 cleavage and repair occurring before the first S-phase would lead to uniform editing, as there is just a single copy of each chromosome. Cas9 activity after the first S-phase increases the likelihood of mosaicism, as there are two or more potential targets on autosomes. Understanding the timing of DSB induction and repair at on-target and off target sites is thus of fundamental importance to evaluate the potential for gene editing in embryos.

Here we use timed collection and timed inhibition using the Anti-CRISPR protein, AcrIIA4, to evaluate the timing of editing at on-target and off-target loci in human embryos. We show that Cas9 editing occurs primarily between 12-16h post-fertilization, corresponding primarily to the G2 phase of the first cell cycle. Cleavage stage embryos contain high levels of mosaicism at both on- and off-target locations, indicating that Cas9 cleavage occurs primarily after S-phase and continues beyond the first cell cycle. Introduction of AcrIIA4 into Cas9 at 16h or ∼8h before mitosis allows for on-target editing but does not prevent chromosomal aneuploidies. Chromosomal consequences of Cas9 cleavage are common throughout the genome, independent of target location, including at MYBPC3. Though this study shows that Cas9 activity is amenable to temporal control, the slow kinetics of Cas9-cleavage and repair relative to the first S-phase remains a limiting factor in reducing embryo mosaicism and preventing aneuploidies.

## RESULTS

### Cas9-induced inDels form primarily after 12h in human embryos

To determine the timing of Cas9-induced edits, Cas9 RNP with sgRNA was injected into donated oocytes at the time of ICSI (**Fig 1A**). We used pools of gRNAs to maximize the number of targets that could be analyzed from a single fertilized embryo. 10 oocytes were thawed along with sperm from normal gamete donors with no known genetic abnormalities. Each egg was injected with Cas9 RNA and three sgRNAs: one targeting the DMD locus on chromosome X, the MYBPC3 locus on chromosome 11, and a guide RNA targeting three sites on the p arm of chromosome 16 and one site on the q arm of chromosome 17, a total of 6 target sites (**Table S1**). These guide RNAs have been characterized in previous studies (7, 8). 9/10 eggs fertilized as indicated by the presence of two pronuclei. Biopsies of the individual haploid pronuclei were harvested at 12 hours (n = 9) or 16 hours (n = 9) post-fertilization. A total of 85/126 targeted sites yielded Sanger genotyping results (12-hour group: 46/63, 16-hour group: 39/63). At 12 hours, corresponding to late S-phase (10), only 11/46 (24%) detectable sites revealed an inDel at the intended cut-site. At 16 hours, corresponding to G2 phase (9), 32/51 (63%) showed an inDel (p = 0.0002, Fisher’s exact test). Four Polar bodies and two cumulus cell samples were also analyzed to serve as a control for genome amplification and Sanger sequencing. In contrast, polar bodies and cumulus cells yielded 47/48 genotyping targets, of which only 3 (7%) were modified (**Fig. 1B**, **Table S2**). This suggests that target amplification was efficient, and editing was inefficient during and after the window of polar body extrusion. Inefficient target amplification of embryo pronuclei may be due to Cas9 cleavage without repair, or amplification failure. Cumulus cells were unmodified for all targets analyzed (0/17) (**Fig. 1B**, **Table S2**).

**Figure 1.**
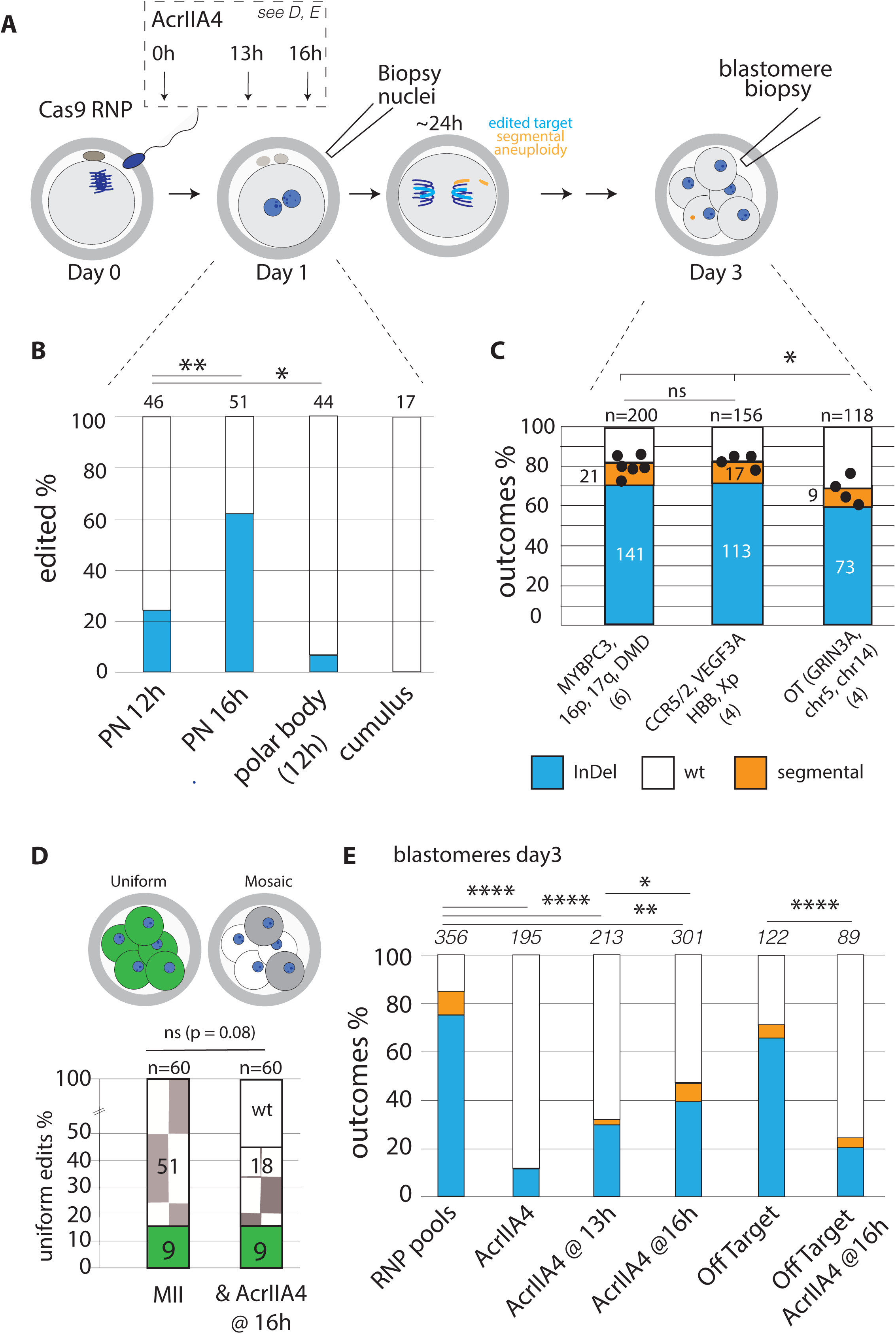
InDel formation requires over half a day, extending into the following cell cycle. **A**) Schematic of the experiment. Human oocytes are fertilized with sperm and a basket of Cas9 RNP targeting different chromosomal locations is applied at fertilization. Individual haploid nuclei are collected at the 1-cell stage, and individual blastomeres at the cleavage stage. In some experiments, the Cas9 inhibitor AcrIIA4 is injected at fertilization together with Cas9 RNPs, or at 13h, or at 16h post fertilization and individual blastomeres collected at the cleavage stage. Cleavage stage blastomeres are analyzed by on-target PCR and Sanger sequencing for InDel formation, and by whole genome high density SNP arrays for chromosomal content. **B**) Editing efficiency indicated with blue bars at target sites. Number of total alleles identified is indicated above each column. **C**) Editing efficiency after 3 days of development of different gRNA RNP baskets, as well as at off-target sites. Blue bars indicate InDels, orange bars indicate segmental aneuploidies. Each dot is the average percent modifications (InDels plus segmental aneuploidies) for each of the different targets. The number of embryos used for analysis is indicated with the Cas9 RNPs. **D**) Mosaicism at specific target sites. Each target site in an individual embryo corresponds to n=1. The total number of embryos analyzed is 8 for each group. AcrIIA4 treated embryos show a lower overall number of edited targets, while the number of uniformly edited targets remains constant. **E**) Editing efficiency and segmental changes depending on indicated condition on day3 of development. The total number of alleles with an editing or chromosomal result is indicated above each bar. **** p<0.0001, ***p<0.001 **P<0.01. * p<0.05. ns = not significant. Fisher’s exact test.

We next compared the editing efficiency of two pools of 3 and 4 guide RNAs, encompassing 6 and 4 on target sites, as well as 4 off-target sites with 2-3 mismatches (**Table S1**). One pool, composed of gRNAs targeting MYBPC3, DMD, and 16p/17q, the same as evaluated at the 1-cell stage. The other pool consisted of gRNAs targeting the Hemoglobin Beta gene (HBB), CCR5, VEGF3A, and a site on chromosome Xp (**Table S1**). The HBB sgRNA also has a known off-target site on Chromosome 9 at the Glutamate Ionotropic Receptor NMDA Type Subunit 3A (GRIN3A) locus (10), and the VEGF3A gRNA has three off-target sites on chromosomes 5 and 14 (11). The two pools showed no significant differences in the frequency of modified alleles on day 3 of embryo development (Average of 81% and 83%, range 73-86%) (**Fig. 1C**). The composition of the modified alleles consisted of 70% and 72% Indels respectively, and 11% segmental changes for both pools. Thus, the two pools are comparable in efficiency, and the variation in efficiency among different guide RNAs is not significant. The four off-target sites with 2-3 mismatches showed modified alleles of 69% on average (range 61-78%), consisting of 61% Indels and 8% segmental errors. This 10% overall reduction at off-target sites compared to on-target sites is surprisingly small, but still significantly different from on-target sites (p<0.01).

Given the late formation of inDels relative to S-phase, replicated sister chromatids may provide separate targets for the same parental chromosome, resulting in mosaicism after cell division. Indeed, in 8 embryos with a total of 60 edited target sites, only 9 sites (15%) showed uniform editing (**Fig. 1D, Table S2**). All embryos were mosaic for at least one of the targets in the gRNA pools. These results show that the majority of Indels emerge more than 12h post injection, and primarily after the first S-phase.

### DSBs lead to segmental aneuploidies even when induced hours before mitosis

In addition to small inDels, chromosomal analysis by SNP arrays revealed segmental chromosomal aneuploidies at all sites, including on chromosomes 3, 5, 6, 9, 11, 14, 16, 17 and chromosome X (**Supplementary** Figure 1). Because chromosomal abnormalities arise in cell division, the timing of DSB induction relative to mitosis may determine whether a DSB can be repaired as an InDel or results in aneuploidy. Potentially, timed inhibition of Cas9 prior to mitosis may allow us to separate editing outcomes from chromosomal consequences. Additionally, this timed inhibition approach may provide insights into the kinetics of DSB repair in human embryos..

To limit Cas9 activity to a specific window, we used the Cas9 inhibitor AcrIIA4 (6). We first determined whether AcrIIA4 protein inhibited Cas9 activity in human embryos. Cas9 RNPs were injected together with AcrIIA4 at the time of fertilization **(Figure 1A)**. As before, we used multiplexing of three unique RNPs to to monitor a larger number or sites (**Table S1**). In 32 blastomeres of 5 embryos, a total of 195 targets were successfully analyzed, of which 22 contained inDels (11% inDels, 89% wild type), significantly lower compared to injections of Cas9 RNPs alone (p=0.00001, **Fig. 1E**). None of the blastomeres contained Cas9-induced segmental aneuploidies. Thus, AcrIIA4 reduces the activity of Cas9 RNP in human embryos by over 7-fold.

We next used AcrIIA4 to inhibit Cas9 activity at two specific time points: at the end of S-phase at 13h post ICSI and RNP application, and in G2 phase, at 16h after ICSI (**Fig. 1A**). A total of 35 blastomeres from 7 embryos were analyzed for the 13h time point and 57 blastomeres from 12 embryos were analyzed for the 16h time point. At both time points, AcrIIA4 reduced the number of edited targets, and increased the percentage of wild type alleles (**Fig. 1E**). Indels were reduced to 30% at the 13h injection time point, and down to 38% at the 16h injection time point, while segmental aneuploidies were at 2% and 8% of alleles, resulting in an overall modified allele percentage of 32% and 46% respectively, a 15% difference within 3 hours. The overall editing efficiency at both time points was significantly lower than without AcrIIA4 (p=0.00001). However, both conditions produced more inDels than when AcrIIA4 was applied at fertilization (p=0.00001). Additionally, editing was more efficient at the 16h time point compared to the 13h time point (p=0.03). In addition to on-target editing, off-target editing was also reduced (**Fig. 1E,** p<0.00001). Thus, the formation of inDels occurs throughout the first cell cycle, and potentially beyond. Furthermore, Cas9 can be inhibited by timed injection of AcrIIA4.

AcrIIA4 increased the frequency of uniform editing but also increased the percentage of unmodified embryos. In the RNP only group, all 60 examined targets showed editing, with only 9 targets (15%) exhibiting uniform editing across all blastomeres of a particular embryo. In contrast, in the AcrIIA4 group, only 18/60 targets showed mosaic editing, and 9 (15%) showed uniform editing (**Fig. 1D, Table S2, Table S3**). Thus, while AcrIIA4 reduces the total number of edited sites, it does not increase the proportion of uniformly edited sites (non-mosaic editing). This suggests that the timing of inDel formation relative to the first S-phase, rather than Cas9 inhibition alone, is the primary factor limiting to preventing mosaicism.

Surprisingly, segmental chromosomal aneuploidies occurred at both time points of Cas9 inhibition (**Fig. 1E**), at 13h (5 segmental errors / 63 inDels, 8%) and the 16h (24 segmental errors / 114 inDels, 21%), not significantly different from RNP only controls (38 segmental errors / 254 inDels, 15%). Mitosis of human zygotes occurs on average at 24h post ICSI (9). Despite inhibition of Cas9 activity 8-12 hours prior to mitosis, DSB ends are not rejoined, and chromosomal aberrations remain common.

### Single alleles at MYPBC3 are primarily due to Cas9-induced chromosomal changes

Cas9-induced chromosomal abnormalities resulting in the loss of the targeted allele and the targeted chromosome have previously been reported for DSBs induced at the EYS and POU5F1 genes. However, other studies proposed that the absence of specific alleles are due to interhomolog repair (12, 13), or due to technical reasons, resulting from allele dropout due to amplification from a single cell (8). While allele dropout may occur when considering a single allele, the absence of heterozygosity across an entire chromosome or chromosome arm seen in SNP arrays is conclusive evidence. Combined with copy number analysis, different outcomes can often be distinguished in single cells. We used identical gRNAs at MYPBC3 as has previously been reported to result in interhomolog repair (8). Of 71 InDels at MYBPC3 detected by PCR and Sanger sequencing, 44 were heterozygous, and 27 cells showed a single allele in Sanger sequencing analysis (**Fig. 2A**, **Table S2**). Of these 27, 3 showed allele dropout in NGS analysis, 1 showed a large deletion of 859bp deleting primer binding sites, 1 had no chromosomal result and was excluded from further analysis, and 19 showed segmental or whole chromosome changes (**Fig. 2A**, **Fig. S2**), Loss of heterozygosity was observed across the p arm, or across the entire chromosome (**Fig. 2B-F**). We also observed failure to amplify the target locus (**Fig. 2E**), which can be caused by complete loss of chromosome segments or an unrepaired DSB (**Fig. 2E**). Chromosomal changes consisted of whole chromosome losses, acentric p arm losses, or p arm gains. Loss of heterozygosity was observed adjacent to the cut site to the telomere, and one cell showed a complete loss of both p arms, and correspondingly no PCR product. The mosaic pattern of inDels and chromosome losses in a 6-cell embryo is consistent with one of the segmental abnormalities occurring in the first cell cycle and a second segmental error in the second cell cycle. The -5bp deletion is a mosaic outcome, though at the time of analysis, chromosomal changes and unrepaired or misrepaired chromosomes made it the only genotype detectable by PCR and Sanger sequencing (**Fig. 2E, F**).

**Figure 2.**
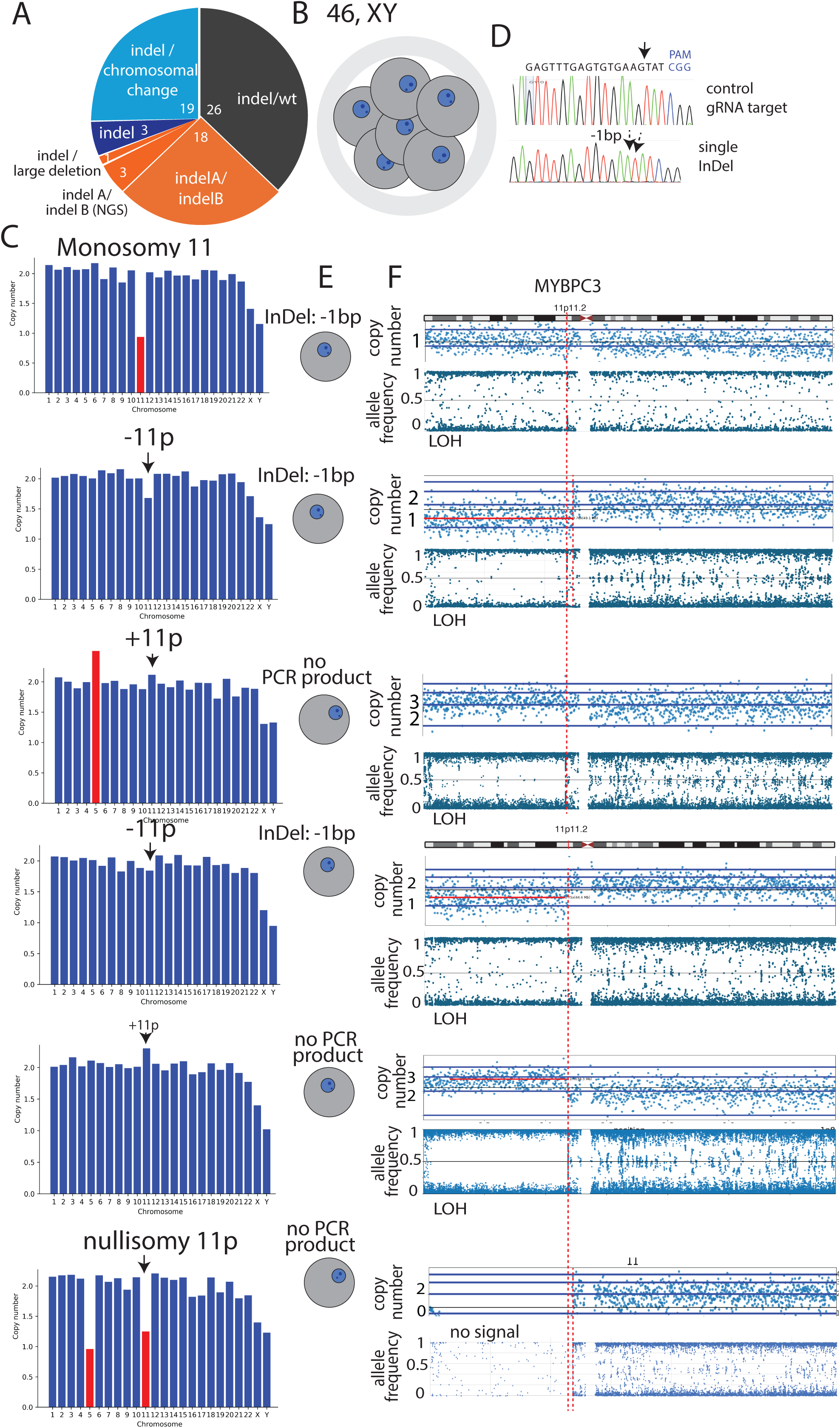
Frequent chromosomal changes due to Cas9 cleavage at the MYPBC3 locus. **A)** Summary of genotypes in cells carrying inDels at MYBPC3. Note that the majority of single inDels are heterozygous with a chromosomal change, not homozygous. IndelA/IndelB are inDels distinguishable by Sanger sequencing. ‘Indel’ is either a single allele or two identical alleles. The number of alleles is indicated. **B**) A male embryo consisting of 6 cells on day 3 was analyzed for chromosome content and InDels at MYBPC3. **C-E)** Each horizontal line are data from a single blastomere. **C**) Chromosome content. SNP array signal intensity is displayed as copy numbers for each chromosome. The cell at the top shows a reduction of copy number of chromosome 11. Two cells show reciprocal mitotic aneuploidy for chromosome 5. **D**) 1bp deletion carried by cells of this embryo by Sanger sequencing. Shown are gRNA and cleavage position. **E**) Sanger sequencing result for each blastomere. 3 blastomeres had no PCR product at the target site, and three showed lack of heterozygosity, with only a single allele carrying a 1bp deletion. **F**) Copy number and heterozygosity detected through B allele frequency on high density SNP arrays. A and B alleles are randomly designated alternate allele identities on the SNP array. A homozygous B allele is displayed as a B allele frequency of 1, a homozygous A allele as a B allele frequency of 0, and heterozygous alleles are at 0.5, with deviations from the mean due to amplification bias. Random noise is likewise in between with random distribution. All cells show loss of heterozygosity across a chromosome arm or the entire chromosome.

Loss of heterozygosity events due to chromosomal changes were also seen at HBB. Hemizygous edited cells showed loss of heterozygosity and copy number changes on either side of Cas9 cleavage (**Fig. S3A-E**). Of 64 edited alleles, 30 were compound heterozygous, 14 were indel/wt, and 18 cells showed a single allele in Sanger analysis. Of these, 3 were heterozygous after NGS analysis, 16 were hemizygous due to chromosomal copy number changes, and one failed chromosomal analysis due to poor quality of amplified DNA and was excluded from analysis (**Fig. S3F, Table S4**). As for the MYBPC3 locus, chromosomal changes at HBB explain the single alleles detected through on target analysis, and others due to allelic bias introduced by WGA from single cells, which masks heterozygosity in a Sanger assay.

At the MYBPC3 locus, there were a small number of apparently homozygous edits (3/70, 4%), without apparent chromosomal copy number changes, deletions of up to 1kb (**Fig. 2A**, **Fig. S3F**). Several different interpretations are possible for the remaining homozygous alleles: hemizygosity due to a deletion or chromosomal rearrangement that is not detected by the PCR assays used or due to an unrepaired DSB. Furthermore, true homozygosity independent identical edits on both chromosomes or due to interhomolog repair. Though independent identical editing at both alleles has a low probability, Indel formation is not random. The most common individual modifications at MYBPC3 in two independent studies were a 1 bp deletion (13%, 18/137 independent events), and a 5bp deletion (12%, 16/137 of independent events), (nonrandom distribution, chi-square p<1e-23, **Table S5**)(8). 1bp deletions can arise from staggered Cas9 cleavage (14), while the recurrent 5bp deletions arise through microhomology mediated end joining, a frequently used pathway in human embryos (4). The probability of two independent identical inDels at MYBPC3 is thus 1.7% and 1.4% respectively, individually low, but cumulatively not insignificant (>3%). Indeed, one of the three apparently homozygous edits was a - 1bp deletion. Preferential repair can thus account for a portion of cells with a single detectable allele on two different chromosomes. At the MYBPC3 and HBB locus combined, only 2/130 edited alleles (1.5%) remain unexplained, with several possible interpretations.

This study establishes that the most common outcomes of a Cas9 induced DSBs are InDels and chromosomal alterations, including segmental and whole chromosome errors, regardless of the chromosomal location. Chromosomal changes were frequent at MYBPC3 and at other sites, regardless of whether Cas9 RNP was applied at MII stage or at the zygote (2PN) stage (**Fig. S1C, Table S2, Table S3**). However, there are distinctions in frequency depending on the site: chromosome loss at MYBPC3 was particularly high: for every segmental error, there are ∼ 2 InDels, significantly higher than at HBB and at DMD (**Fig. S1B**). The molecular mechanisms underlying preferences in DSB repair outcomes depending on the genomic location are not currently known.

## DISCUSSION

The integrity of the genome is a primary determinant of developmental potential. Cas9-mediated gene editing provides a tool to interrogate DNA repair outcomes in the early embryo. These studies reveal that a DSB results in frequent chromosomal changes, and that the repair is slow and inefficient.

Several studies have documented the slow kinetics of Cas9-induced DSB repair in other cell types. In somatic cells, across different loci, the half-life of a DSB until repair ranges from 1.4-10.7 hours (15). Formation of Indels was found to require 10-30 hours, with repair by microhomology-mediated end joining delayed by ∼10 hours relative to nonhomologous end joining. MMEJ, also termed Alt-EJ, is a repair pathway involving resection of DNA ends and is a common repair outcome in both human embryos and human embryonic stem cells (4). NHEJ may result in re-joining without alteration of an unprocessed site, followed by recurrent cleavage, the kinetics of which cannot be detected in the assays used here. However, the processing and repair of DSBs resulting in a sequence alteration that is no longer a match to the gRNA requires many hours in human embryos. Similarly, in iPS cells, InDels form slowly, within days, rather than hours (16). In neurons, DSBs can remain unrepaired for weeks, with InDels forming extremely slowly. Consistent with the findings in pluripotent stem cells, we show that InDels begin to form in human embryos around 12 hours post-fertilization, increasing to 40-60% of targets by the 16-hour mark, and continuing to day 3 to reach an overall editing efficiency of >80%. This indicates that inDels form predominantly after the first DNA replication cycle, which is largely complete by 12h post fertilization (9). The slow kinetics of DSB repair and InDels formation is not unique to the human embryo, but the consequences are distinct. In somatic cells, a more effective response to a DSB causes G1 arrest and delays the progression to mitosis (17). When checkpoints are experimentally inhibited in somatic cells, chromosomal abnormalities and micronucleation follow (17). In human embryos, checkpoints are intrinsically weak, and one, or even several spontaneous or Cas9-induced breaks fail to prevent entry into mitosis, resulting in aneuploidy and micronucleation (4, 9).

Across more than ten Cas9 target sites examined, we found frequent segmental chromosomal abnormalities consistent with unrepaired Cas9-induced breaks. Whole chromosome aneuploidies are also a consequence of Cas9 cleavage (4). While we find whole chromosomal changes on targeted chromosomes, we did not systematically examine this aspect here, as the gRNAs used are not allele-specific and thus spontaneous and Cas9-induced whole chromosome changes cannot conclusively be distinguished. Interestingly, different Cas9 target sites show varying frequencies of segmental chromosomal changes, with MYBPC3 showing particularly frequent aneuploidies. The mechanisms determining these differences are not currently known but may be related to differences in kinetics at different DSB repair sites, which varied from ∼1-10 hours in cultured human CML cells (15).

The slow kinetics of Cas9-induced InDel formation has two major consequences: embryo mosaicism and chromosomal aneuploidy. These two consequences are not readily separable: when Cas9 is inhibited using AcrIIA4 hours before the first mitosis, indel efficiency is reduced, but aneuploidies still occur, despite 8-12 hours available for repair prior to cell division. Cas9 has been approved for somatic gene therapy for sickle cell disease and has been considered for germ line gene therapy as well, with at least one unpublished report of the use of Cas9 to delete CCR5 to increase HIV resistance (18). The loss of a targeted mutation has previously been interpreted as evidence for correction and efficient repair between homologous chromosomes at MYBPC3 (12), which appeared to elevate such promise. However, at that the time of these studies, the chromosomal consequences of Cas9 had not been known, and both clinical progress and interpretation of results assumed a binary outcome of a DSB, of repair by homologous recombination or InDels. This emphasizes the critical importance of basic research agnostic to clinical relevance, with a broad view toward identifying unknown consequences of gene editors. The frequent chromosomal changes at MYBPC3 reported here, alongside large deletions, and other difficult-to-detect genetic changes, including unrepaired DSBs, leave little room for interhomolog repair as a possible interpretation of the frequent loss of a targeted mutation. Based on other studies in non-meiotic cells, interhomolog recombination resulting in copy-neutral loss of heterozygosity is expected to be rare (19, 20).

Double strand breaks are among the most toxic lesions in the genome because a single break can potentially be cell lethal, in particular in dividing cells. Due to slow kinetics of repair, causing mosaicism and frequent chromosomal abnormalities, as well as large deletions Cas9 is an implausible candidate for the clinical correction of mutations in human embryos. To a lesser extent, this may also be relevant to more differentiated cells, where rearrangements, micronucleation, and chromosome-scale losses have also been observed (19–23), though not at the staggeringly high frequency seen in human embryos. Regardless of the challenges to use in gene editing, there is great utility for Cas9 and other gene editors to study DNA repair and cell cycle checkpoint control as a determinant of developmental potential.

## METHODS

The Columbia University Institutional Review Board approved all human procedures and experiments, including the use of donated oocytes and sperm for fertilization, Cas9 to study DNA repair in human embryos, and the sharing of array data.

### Gametes

Cryopreserved human oocytes, semen samples, and zygotes were anonymously donated from individuals who provided informed consent for use in research. Oocytes were cryopreserved between the years 2012 and 2014 using Cryotech vitrification kit. Sperm samples were cryopreserved between the years 2016 and 2018 using Quinn’s washing medium from Cooper Surgical and TYB freezing media from Irvine Scientific. All gametes were stored in liquid nitrogen until use.

### Protein expression and purification

A bacterial culture harboring the plasmid #101043 with GST-AcrIIA4 fusion protein was obtained from Addgene. The bacteria were streaked on LB Agar plates with Ampicillin for overnight incubation at 37°C. Individual colonies were then isolated and resuspended in Luria Broth (LB) with Ampicillin and incubated in a shaking incubator at 37°C for 18 hours. The bacteria were then diluted 1:100 in LB medium with ampicillin and incubated in a shaking incubator at 37°C until an Optical Density of 0.4-0.6 was reached. The GST-AcrIIA4 fusion protein was then overexpressed with overnight induction of 0.25 mM isopropyl-β-D-thiogalactopyranoside (Research Products International # I56000-1.0) at 20°C in a shaking incubator. The culture was centrifuged, and the supernatant was discarded. The bacterial pellet was resuspended in PBS with 1% Triton and the bacterial cells were lysed using sonication. The fusion protein was then purified by centrifugation followed by the addition of Glutathione Sepharose 4B resin (Cytivia # 17-0756-01) to the supernatant. The recovered proteins were washed with PBS three times and centrifuged, keeping only the pellet. The pellet was then washed with cleavage buffer (50 mM Tris-HCl, pH 7.0 (at 25 °C), 150 mM NaCl, 1 mM EDTA, 1 mM dithiothreitol) and digested with PreScission Protease (GenScript Z02799-100) to remove the GST tag by chilling the mixture at 5°C C for 4 hours and separating with centrifugation once more. The purified AcrIIA4 protein was stored in cleavage buffer (50mM Tris-HCl (pH 7.0), 150 mM NaCl, 1 mM EDTA, and 1 mM dithiothreitol (DTT)) and frozen in multiple aliquots at −20°C.

### Protein purification confirmation

Isolated AcrIIa4 protein was run on sodium dodecyl sulfate-polyacrylamide gel electrophoresis (SDS-PAGE) with Coomassie blue staining to confirm protein size **(Supplemental Figure 3).** The purified protein was also confirmed using Western Blot with an AcrIIA4 primary antibody and anti-mouse secondary antibody (Cell Signaling Technology # 7076S) **(Supplemental Figure 3).** Protein concentration was determined through a Bicinchoninic acid assay.

### RNP preparation

2 nmol of single guide RNAs were obtained from Integrated DNA Technologies (IDT). Individual gRNA sequences are listed in Supplementary Table 1. Each gRNA was dissolved in 20 ul of nuclease-free water to make a 100 <u>μ</u>M sgRNA concentration. Ribonucleoprotein (RNP) preparation was performed as previously described (Turocy, 2022). Briefly, 3 μL of injection buffer (5mM Tris-HCl, 0.1mM EDTA, pH 7.8), 2 μL of 63M IDT nlsCas9 v3, and 1.5 μL of 100M sgRNA were combined and incubated at room temperature for 5 minutes. After, 96.5 μL of injection buffer was added to the mix. The solution was then centrifuged at 16000 RCF for 2 minutes and then placed on ice prior to intracytoplasmic sperm injection (ICSI).

1.5 nmol of single guide RNA were obtained from Synthego. Individual gRNA sequences are listed in Supplementary Table 1. Each gRNA was dissolved in 15 μL of nuclease-free 1X TE buffer per manufacturer instructions to a concentration of 100 μM. RNP preparation involved combining 3.125 μL of 20 μM Cas9 V3 (New England Biolabs), 0.776 μL of 100 μM sgRNA and incubation at room temperature for 10 minutes. Thereafter, 46 μL of injection buffer was added. The solution was then centrifuged at 16000 RCF for 2 minutes and placed on ice prior to ICSI.

### Oocyte and Zygote Manipulation

Oocyte manipulations were performed in an inverted Olympus IX71 microscope using Narishige micromanipulators. The platform was heated to 37°C. Frozen oocytes were thawed using Cryotech Warming solution set 205. Cryopreserved sperm were thawed to room temperature for 10 min. Quinn’s Sperm Washing Medium was added to the thawed sperm and centrifuged at 300x g for 20 min. A second wash was performed and centrifuged again. The sperm pellet was then suspended in 200 μL of wash media and analyzed for viability. Manipulation dishes consisted of a droplet with Cas9 RNP with or without 12.5 μM of AcrIIA4 diluted in injection buffer. Individual motile sperm were immobilized by pressing the sperm tail with the ICSI micropipette, picked up, and expelled in the Cas9 RNP droplet.

The sperm was then picked up again for injection into the oocyte cytoplasm. All eggs were cultured in Global total media in an incubator at 37°C and 5% CO2. Pronucleus formation to confirm successful fertilization was done on day 1 after ICSI. Experiments with a delayed injection of ACrIIa4 were performed between 16-17 h post-ICSI. The tip of an injection needle was nicked, and small amounts of the AcrIIa4 diluted in injection buffer was injected manually into the oocyte cytoplasm using a Narishige micromanipulator.

### Genome amplification and Genotyping

Zygotes were collected at 12 or 16 h post-ICSI, and single blastomeres were collected on day 3 using the inverted Olympus IX71 microscope. Narishige micromanipulators and a zona pellucida laser (Hamilton-Thorne) were used to aid in the collection of the blastomeres. All samples were placed in PCR tubes containing 2 or 4 μL of PBS for half of full reactions. REPLI-g single kit (Qiagen) which was used for amplification according to the manufacturer’s instructions, with either a half-reaction of 25 μL or a full reaction of 50 μL. Genotyping was performed with custom primers flanking each cut-site as listed in Table S5.

PCR was performed using AmpliTaq Gold, and all products were run on a 1.5% agarose gel for visual inspection. All samples with a visible band were submitted to Azenta (Genewiz) for Sanger sequencing. Cas9-mediated gene editing was analyzed using NCBI Blast reports and ICE analysis (Synthego).

### Amplicon next-generation sequencing

Indicated DNA samples were amplified using their respective primers (**Supplemental Table S6**). Each sample underwent a precipitation reaction to purify the DNA. Briefly 1/10 volume of 3M sodium acetate was added to PCR products, followed by 3x volume of 100% ethanol. The solution was mixed and incubated at -20°C for 20 minutes. The sample was centrifuged at 18000G for 20 minutes as room temperature and the supernatant dumped. The pellet was then washed with 800 μL of 70% Ethanol and centrifuged at 15000g x 20 minutes at room temperature. The supernatant was removed and the pellet was allowed to air dry on a hot plate for 45 minutes. The pellet was reconstituted in 20 μL of nuclease free water. Concentration was checked with nanodrop and normalized to 20 ng/μL. Samples were sent to Azenta for AmpliconEZ analysis. Analysis of variant frequency was performed by Genewiz and by CRISPResso2.

### Genome-wide SNP array

Genome-wide SNP array was performed as previously described (Zuccaro et al., 2020). Briefly, embryo and blastomere biopsies were amplified using REPLI-g single cell kit according to the manufacturer’s instructions or at Genomic Prediction using ePGT amplification as previously described (24), or using the Qiagen REPLI-g Single Cell Kit (Cat #150343). Copy number and genotyping analysis were performed using gSUITE software (Genomic Prediction). To determine copy number, raw intensities from the Affymetrix Axiom arrays, which contain data from 820,967 SNPs across all chromosomes, were processed and subsequently normalized with a panel of wild type male and female samples.

Chromosomal breakpoints were mapped by identifying break sites based on signal intensity of probes, as well as transitions in heterozygosity. For samples with 1 to 2 copies difference, genomic coordinates were determined graphically based on copy number plots. A segmental error was defined as the gain or loss of a chromosome segment centromeric or telomeric to the Cas9 cut site coordinate. For loss of heterozygosity analysis, background was removed by excluding probes with negative Log2P values. Data are available at GEO accession number GSE288845.

### Quantification and Statistical Analysis

Statistical analysis was performed using Fisher’s exact test. Non-random distribution of inDel size was tested using Chi-square test.

## AUTHOR CONTRIBUTIONS

AR and DE designed the study. AR performed AcrIIA4 expression and isolation, Western blotting, sperm preparation, RNP preparation, genotyping, chromosomal analysis and data interpretation, with contributions from SJ. DM, JX and NT performed SNP array analysis. JT performed PCR and genotyping, including on target NGS. JS performed analysis of chromosomal break site positions. DE performed embryology. AR and DE wrote the paper.

## ACKNOWLEDGMENTS

This work was supported by the US-Israel Binational Science Foundation *#2021278*. A.R. was supported by an ObGyn research fellowship. S.J. was supported by a NYSCF-Druckenmiller postdoctoral fellowship. GP provided all SNP array analysis. Embryos and gametes were donated.

---

**Supplemental Figure 1.**
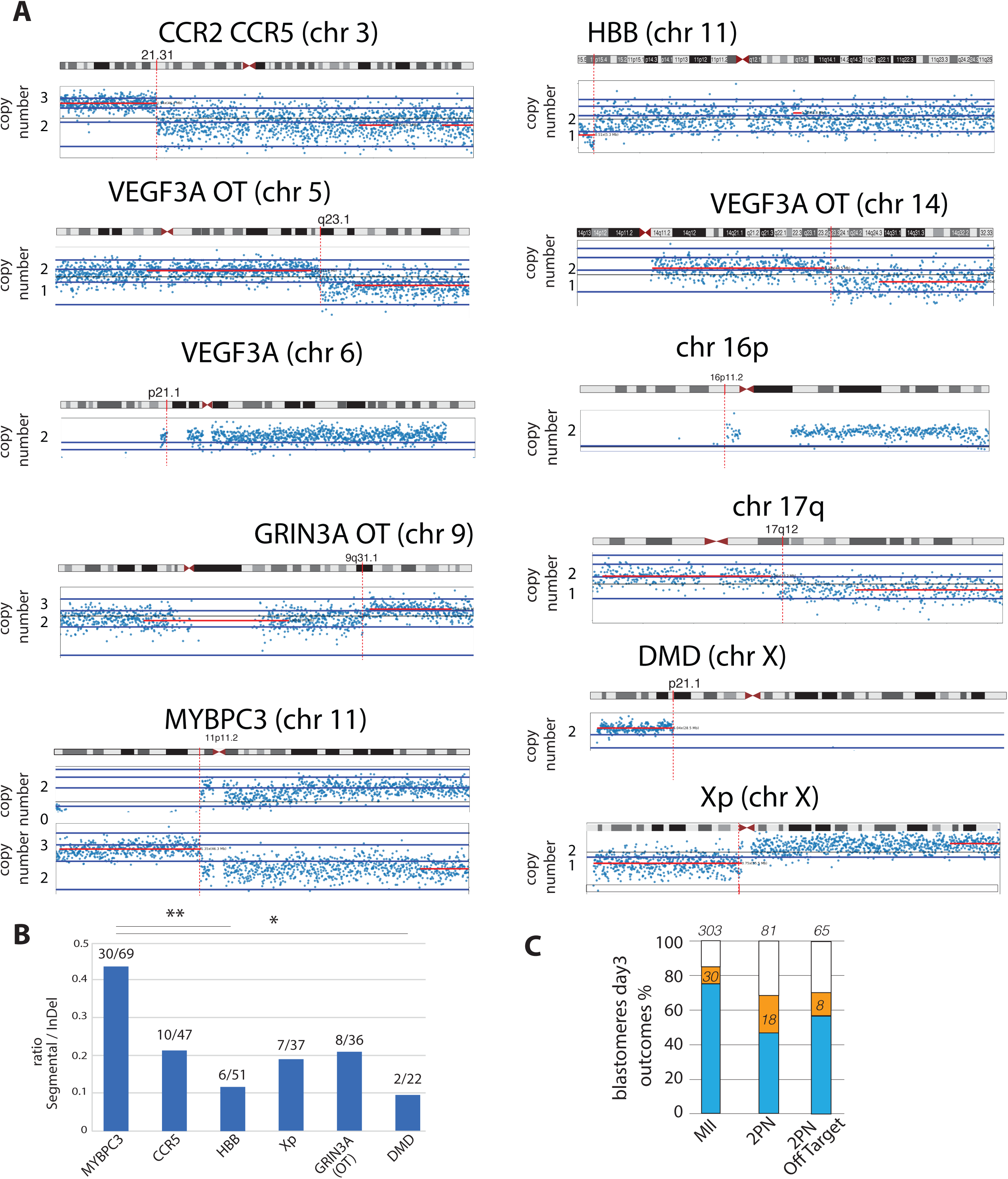
Segmental aneuploidies at 11 target sites throughout the genome. **A)** Shown are SNP array signal intensities for specific chromosomes, reflecting copy number. Cas9 gRNA target sites are shown with a red line on the respective chromosomes. **B)** Frequency of segmental aneuploidies relative to InDels. Shown are the number of cells with segmental aneuploidies and the number of detectable InDels alleles. Nullisomic aneuploidies or heterozygous segmental aneuploidies were both counted as a single chromosomal change. The number of whole chromosome changes induced by Cas9 is not evaluated, because the targeting is not allele specific, and cannot conclusively be distinguished from spontaneous aneuploidies. Thus, the number of chromosomal changes per target is a conservative count across all sites. ** p<0.01, * p<0.05. Fisher’s exact test. **C)** Chromosomal changes when baskets of RNPs are applied at the MII stage or at the 2PN stage on day1 of development. The number of genotyped total alleles is provided on top of each bar. Related to Fig. 1.

**Supplemental Figure 2.**
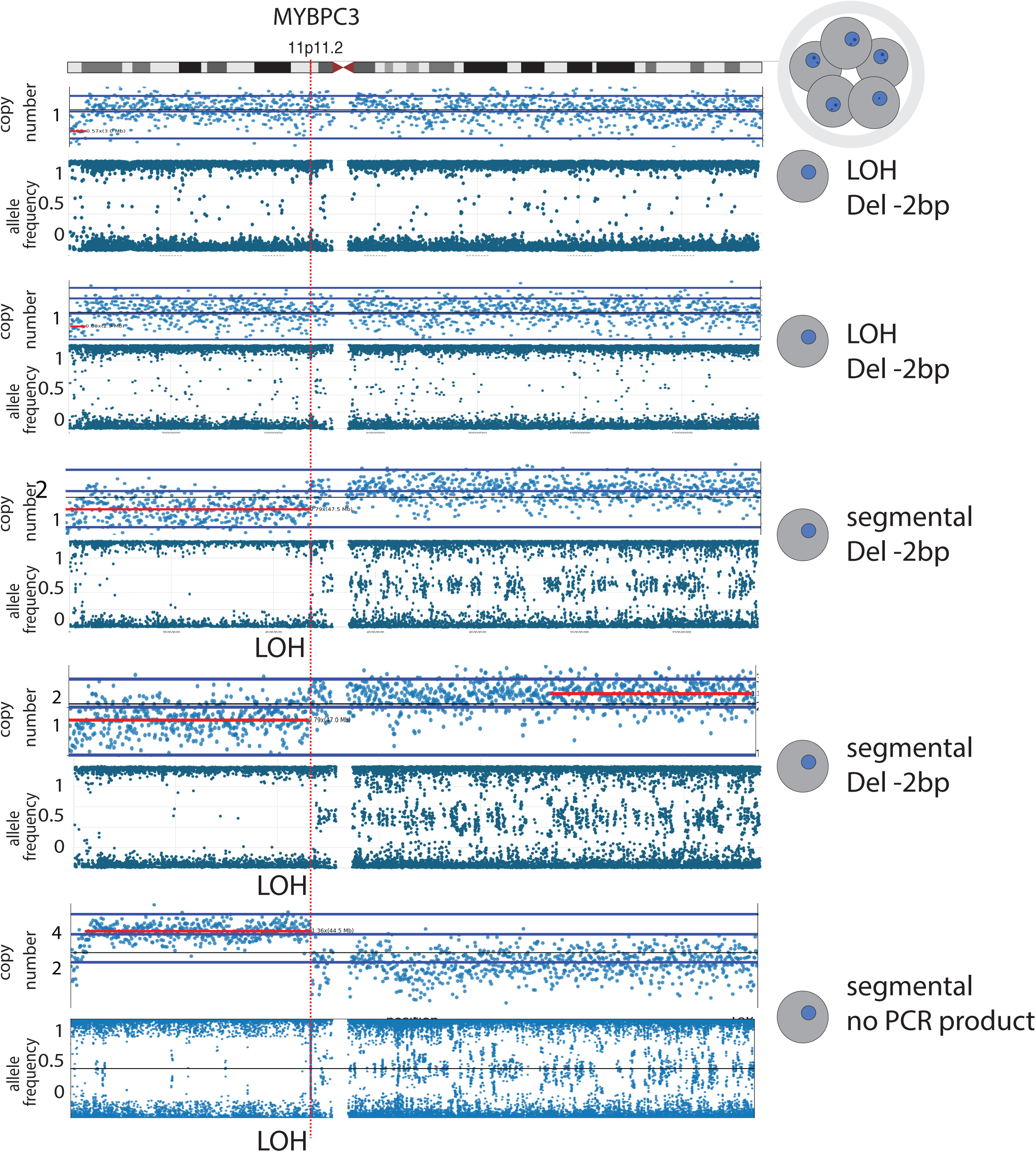
Cas9-induced chromosomal changes at the MYPBC3 locus. Shown are chromosomal analysis by SNP array, copy number and allele frequency, as well as on target Sanger sequencing results of a 5-cell embryo on day3 of development. Note the loss of heterozygosity either across the entire chromosome or on chromosome 11p for all cells. Related to Fig. 2.

**Supplemental Figure 3.**
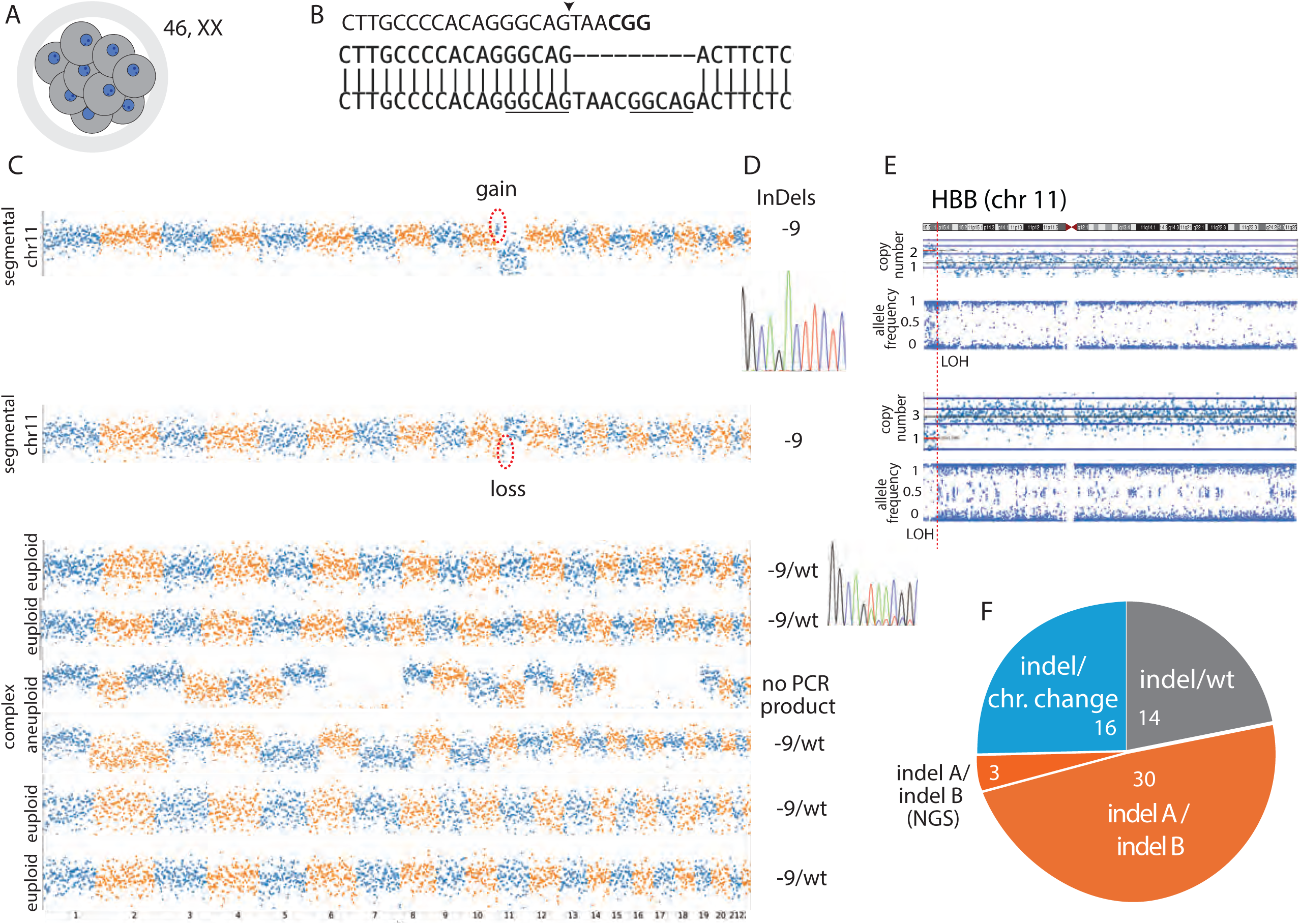
Cas9-induced chromosomal changes at the HBB locus. **A)** Schematic of the embryo analyzed. **B**) gRNA target site. PAM is bold, cut site is indicated with an arrowhead, and microhomology is underlined. Shown is the wt reference sequence and the -9bp deletion. **C**) Shown are copy number plots across the entire genome for 8 cells of a 9-cell embryo on day3 of development. Segmental aneuploidy on chromosome 11 is circled. These two cells show only a single edited allele by PCR and Sanger sequencing (**D**). Copy number change and LOH from the Cas9 cleavage site through the p or the q arm all the way to the telomere as analyzed by SNP array (**E**). Other cells show heterozygosity for the same 9 bp deletion, and a wild type chromosome (C). Two cells show complex aneuploidy. **F**) Summary of allelic outcomes. The number of alleles is indicated. Note that most homozygous appearing alleles carry chromosomal changes. IndelA and IndelB are distinguishable by Sanger sequencing. ‘Indel’ may constitute a single detectable allele, or two identical alleles. Related to Fig. 2.

**Supplemental Figure 4.**
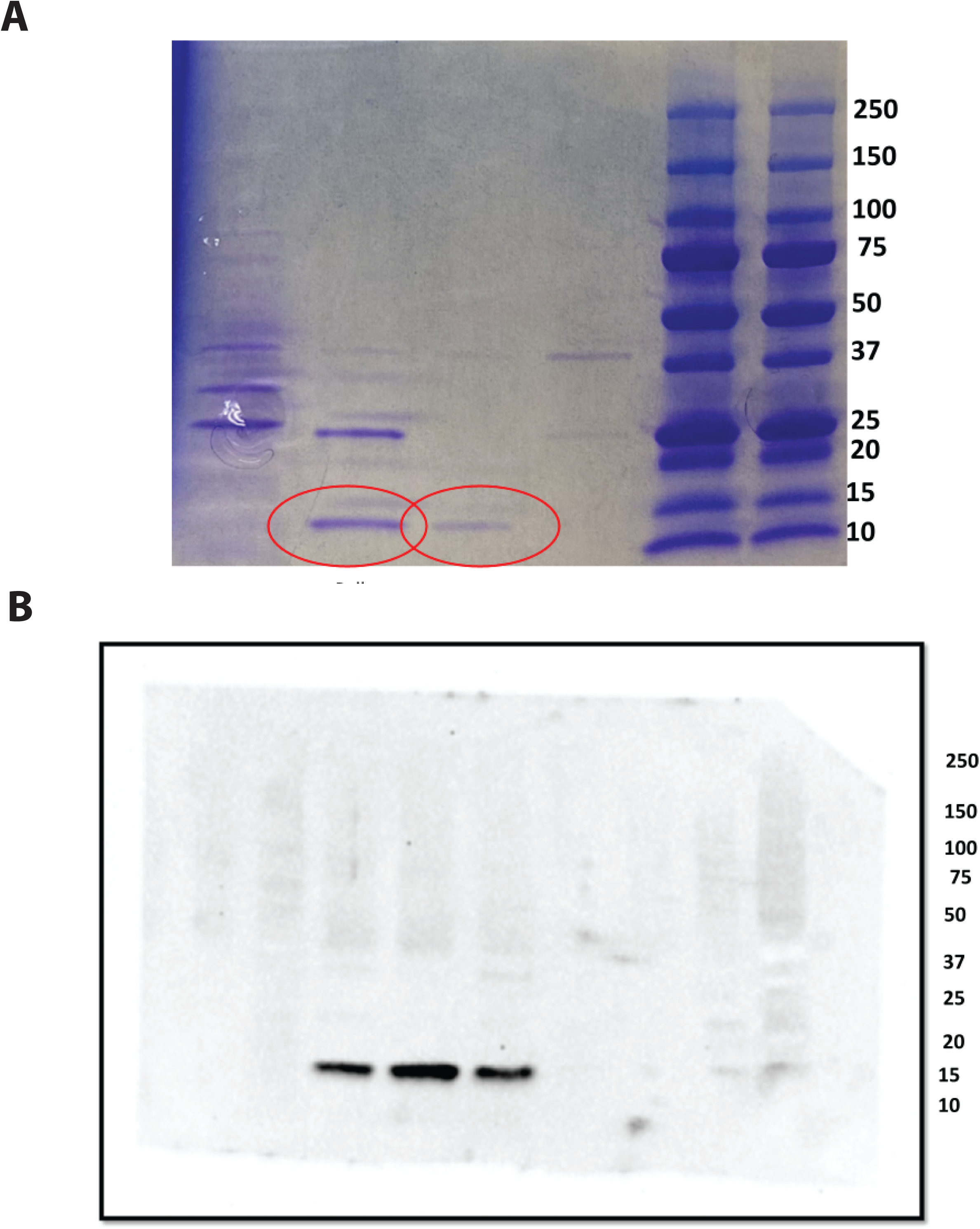
SDS Page with Coomassie Blue Stain for AcrIIA4 protein. A) Image of SDS Page with Coomassie Blue showing AcrIIA4 protein aligned with 10-15 kDa ladder. MW of AcrIIA4 ∼10 kDA. AcrIIA4-GST fusion protein also shown aligned with the 25 kDA ladder. MW of GST ∼15 kDa. B) Western blot of AcrIIA4 with secondary Anti mouse IgG, HRP linked Antibody

**Table S1. gRNA targets and sequences**

**Table S2 Comprehensive list of outcomes for each blastomere and analysis**

Different columns indicate conditions, RNP baskets, use of AcrIIA4, InDels and chromosomal outcomes for each target listed. Columns without a listed result for a specific experiment do not apply because the Cas9 RNP was not used. Colored cells without an entry had no genotyping result.

**Table S3 Summary of Table S2.** This table adds up number of Table S2 at different sites. Related to Fig. 1, Fig. 2A, Fig. S1B, Fig. S1C, Fig. S3

**Table S4 On target deep sequencing analysis of cells with monoallelic Sanger result**

Single molecule sequencing reveals uneven amplification of a second allele, which appears as allele dropout in Sanger sequencing. In contrast, cells with chromosomal changes, show a single allele in both Sanger and on target deep sequencing. Shown are the number of reads detected per allele.

**Table S5 InDels at MYPBC3 are non random.** Number of independent inDels occurring in different embryos. MMEJ contributes to the most common deletions except for the deletion of -1bp.

**Table S6 Primer sequences used for genotyping**

## REFERENCES

1. Rodríguez-Rodríguez DR, Ramírez-Solís R, Garza-Elizondo MA, Garza-Rodríguez ML, Barrera-Saldaña HA. Genome editing: A perspective on the application of CRISPR/Cas9 to study human diseases (Review). Int J Mol Med. 2019;43(4):1559–74.

2. Turocy J, Adashi EY, Egli D. Heritable human genome editing: Research progress, ethical considerations, and hurdles to clinical practice. Cell. 2021;184(6):1561–74.

3. National Academies of Sciences E, Medicine, National Academy of M, National Academy of S, Committee on Human Gene Editing: Scientific M, Ethical C. Human Genome Editing: Science, Ethics, and Governance. Washington (DC): National Academies Press (US) Copyright 2017 by the National Academy of Sciences. All rights reserved.; 2017.

4. Zuccaro MV, Xu J, Mitchell C, Marin D, Zimmerman R, Rana B, et al. Allele-Specific Chromosome Removal after Cas9 Cleavage in Human Embryos. Cell. 2020;183(6):1650–64.e15.

5. Alanis-Lobato G, Zohren J, McCarthy A, Fogarty NME, Kubikova N, Hardman E, et al. Frequent loss of heterozygosity in CRISPR-Cas9-edited early human embryos. Proc Natl Acad Sci U S A. 2021;118(22).

6. Shin J, Jiang F, Liu J-J, Bray NL, Rauch BJ, Baik SH, et al. Disabling Cas9 by an anti-CRISPR DNA mimic. Science Advances. 2017;3(7):e1701620.

7. Turocy J, Marin D, Xu S, Xu J, Robles A, Treff N, et al. DNA Double Strand Breaks cause chromosome loss through sister chromatid tethering in human embryos. bioRxiv. 2022:2022.03.10.483502.

8. Liang D, Mikhalchenko A, Ma H, Marti Gutierrez N, Chen T, Lee Y, et al. Limitations of gene editing assessments in human preimplantation embryos. Nat Commun. 2023;14(1):1219.

9. Palmerola KL, Amrane S, De Los Angeles A, Xu S, Wang N, de Pinho J, et al. Replication stress impairs chromosome segregation and preimplantation development in human embryos. Cell. 2022;185(16):2988–3007.e20.

10. Cradick TJ, Fine EJ, Antico CJ, Bao G. CRISPR/Cas9 systems targeting β-globin and CCR5 genes have substantial off-target activity. Nucleic Acids Res. 2013;41(20):9584–92.

11. Tsai SQ, Zheng Z, Nguyen NT, Liebers M, Topkar VV, Thapar V, et al. GUIDE-seq enables genome-wide profiling of off-target cleavage by CRISPR-Cas nucleases. Nat Biotechnol. 2015;33(2):187–97.

12. Ma H, Marti-Gutierrez N, Park SW, Wu J, Lee Y, Suzuki K, et al. Correction of a pathogenic gene mutation in human embryos. Nature. 2017;548(7668):413-9.

13. Ma H, Marti-Gutierrez N, Park S-W, Wu J, Hayama T, Darby H, et al. Ma et al. reply. Nature. 2018;560(7717):E10–E23.

14. Lemos BR, Kaplan AC, Bae JE, Ferrazzoli AE, Kuo J, Anand RP, et al. CRISPR/Cas9 cleavages in budding yeast reveal templated insertions and strand-specific insertion/deletion profiles. Proc Natl Acad Sci U S A. 2018;115(9):E2040–e7.

15. Brinkman EK, Chen T, de Haas M, Holland HA, Akhtar W, van Steensel B. Kinetics and Fidelity of the Repair of Cas9-Induced Double-Strand DNA Breaks. Mol Cell. 2018;70(5):801–13.e6.

16. Ramadoss GN, Namaganda SJ, Hamilton JR, Sharma R, Chow KG, Macklin BL, et al. Neuronal DNA repair reveals strategies to influence CRISPR editing outcomes. bioRxiv. 2024.

17. van den Berg J, G. Manjón A, Kielbassa K, Feringa FM, Freire R, Medema René H. A limited number of double-strand DNA breaks is sufficient to delay cell cycle progression. Nucleic Acids Research. 2018;46(19):10132–44.

18. Regelado A. Exclusive: Chinese Scientists Are Creating Crispr Babies. MIT Technology Review. 2018.

19. Regan SB, Medhi D, White TB, Jiang YZ, Jia S, Deng Q, et al. Megabase-scale loss of heterozygosity provoked by CRISPR-Cas9 DNA double-strand breaks. bioRxiv. 2024.

20. Boutin J, Rosier J, Cappellen D, Prat F, Toutain J, Pennamen P, et al. CRISPR-Cas9 globin editing can induce megabase-scale copy-neutral losses of heterozygosity in hematopoietic cells. Nat Commun. 2021;12(1):4922.

21. Kosicki M, Tomberg K, Bradley A. Repair of double-strand breaks induced by CRISPR-Cas9 leads to large deletions and complex rearrangements. Nature Biotechnology. 2018;36(8):765–71.

22. Leibowitz ML, Papathanasiou S, Doerfler PA, Blaine LJ, Sun L, Yao Y, et al. Chromothripsis as an on-target consequence of CRISPR-Cas9 genome editing. Nat Genet. 2021;53(6):895–905.

23. Cullot G, Boutin J, Toutain J, Prat F, Pennamen P, Rooryck C, et al. CRISPR-Cas9 genome editing induces megabase-scale chromosomal truncations. Nature Communications. 2019;10(1):1136.

24. Treff NR, Zimmerman R, Bechor E, Hsu J, Rana B, Jensen J, et al. Validation of concurrent preimplantation genetic testing for polygenic and monogenic disorders, structural rearrangements, and whole and segmental chromosome aneuploidy with a single universal platform. Eur J Med Genet. 2019;62(8):103647.

